# PPM1B utilizes a trinuclear metal architecture for phosphatase activity

**DOI:** 10.64898/2026.04.23.720145

**Authors:** Reece P. Stevens, Viktoriya Solodushko, Andrzej Wierzbicki, Thomas C. Rich, Mikael F. Alexeyev, Marlo K. Thompson, Madeline Stone, Camryn Hall, Althea deWeever, Sarah L. Sayner, Troy Stevens, Joel Andrews, Aishwarya Prakash, Richard E. Honkanen, Ji Young Lee, E. Alan Salter, Mark R. Swingle

## Abstract

The metal-dependent protein phosphatase (PPM/PP2C) family regulates innate immune and cell death pathways through reversible phosphorylation. Although these enzymes contain a conserved third Mg^2+^/Mn^2+^ ion (M3) that is essential for activity, its chemical role in phosphate hydrolysis has remained unclear. Here, we report studies that reveal PPM1B promotes cell death during *Pseudomonas aeruginosa* infection and utilizes a trinuclear metal center in which M3 directly coordinates the substrate phosphate, positioning it for in-line S_N_2 hydrolysis. In addition to substrate orientation, M3 positions a water molecule to protonate the departing alkoxide, stabilizing the leaving-group. Functionally, M3 substitutes for the arginine clamp in phosphoprotein phosphatases (PPP), revealing that these evolutionarily distinct phosphatase families have converged on the same chemical strategy through fundamentally different catalytic architectures. Together, these findings define a three-metal mechanism in PPM phosphatases and identify the M3 site as a rare and potentially druggable feature for immune and infectious diseases.

---

Innate immune signaling relies on reversible protein phosphorylation to control the timing and magnitude of host defense responses^1–8^. Although kinase-driven pathway activation has been extensively characterized, far less is known about how phosphatases execute signal termination. PPM1B is a member of the metal-dependent protein phosphatase (PPM/ PP2C) family that negatively regulates key innate immune signaling pathways, including NF-κB, MAPK, cGAS– STING, and RIG-I^1–8^. Despite this broad regulatory scope, its function during bacterial infection has not been explored. Here, we first establish that PPM1B promotes cell death when challenged with *Pseudomonas aeruginosa*, suggesting it has a functional role in dictating cell fate during infection.

These findings prompted us to investigate the catalytic mechanism underlying PPM1B function and the PPM family in general. Notably, the mechanism of action of the PPM family remains poorly defined, in part because these phosphatases lack the canonical Arg clamp used by other phosphatase families to position the phosphate for S_N_2 hydrolysis^9–12^. Recent structural studies on PPM1A and related PP2C homologs have identified a third metal ion (M3) within the active site, coordinated by conserved Asp residues^13–17^. Although disruption of M3 coordination markedly reduces or abolishes phosphatase activity^13–17^, existing models do not explain how M3 contributes to phosphate hydrolysis at the chemical level, nor how catalysis proceeds in the absence of the canonical Arg clamp. As a result, the fundamental catalytic logic of PPM phosphatases remains unresolved.

We hypothesized that M3 directly coordinates the substrate phosphate to enable catalysis, functionally substituting for the Arg clamp. Combining biochemical, structural, and computational approaches, we demonstrate that M3 is essential for catalytic activity but dispensable for substrate binding. Mechanistically, PPM1B assembles a trinuclear metal center in which M3 directly coordinates a phosphoryl oxygen to orient the substrate for in-line nucleophilic attack while also positioning a water molecule for protonation of the exiting alkoxide. Structural comparison with PP5 indicates that M3 performs a role analogous to the conserved Arg clamp used by the phosphoprotein phosphatase (PPP) family to position the phosphate for hydrolysis. We further show that mutation of a conserved M3 coordinating residue (PPM1B^D151E^) causes loss of catalytic activity by disrupting M3 contact with the phosphoryl oxygen and misaligning the substrate phosphate in a nonreactive position. Together, these findings establish a link between PPM1B function in controlling cell fate during infection and demonstrate convergent evolution between the PPM and PPP families, which achieve the same catalytic solution to phosphate positioning through distinct molecular strategies, using either a coordinated metal ion or basic amino acid side chains to enable efficient phosphoryl hydrolysis.

## Results

### PPM1B promotes cell death during *Pseudomonas aeruginosa* infection

We first investigated the role of PPM1B in the host cell response to *Pseudomonas aeruginosa* infection (strain PA103). CRISPR-Cas9 was used to knockout (KO) PPM1B from A549 cells. DNA sequencing showed the introduction of premature stop codons in candidate KO cells, which was accompanied by a significant reduction in messenger RNA when assessed by quantitative PCR (Supplementary Fig. 1). Additionally, PPM1B protein could not be detected by western blot in candidate KO cells (Fig. 1a), demonstrating successful KO of the protein. Wild type (WT) and KO cells were infected with PA103 for 2 hours, followed by assessment of lactate dehydrogenase (LDH) release and LiveDead staining. When challenged with *Pseudomonas aeruginosa*, WT cells had increased media LDH and higher red staining than KO cells (Fig. 1b-d), indicating worse injury in WT cells. To independently validate this observation, we reduced PPM1B expression in WT cells using siRNA, achieving approximately 80% knockdown (Fig. 1e-f). During infection, cells treated with scramble control siRNA had higher LDH release than PPM1B knockdown cells (Fig. 1g), showing consistent results with the KO cells. Overall, our findings suggest that PPM1B promotes cell death in A549 cells during *Pseudomonas aeruginosa* infection.

**Fig. 1.**
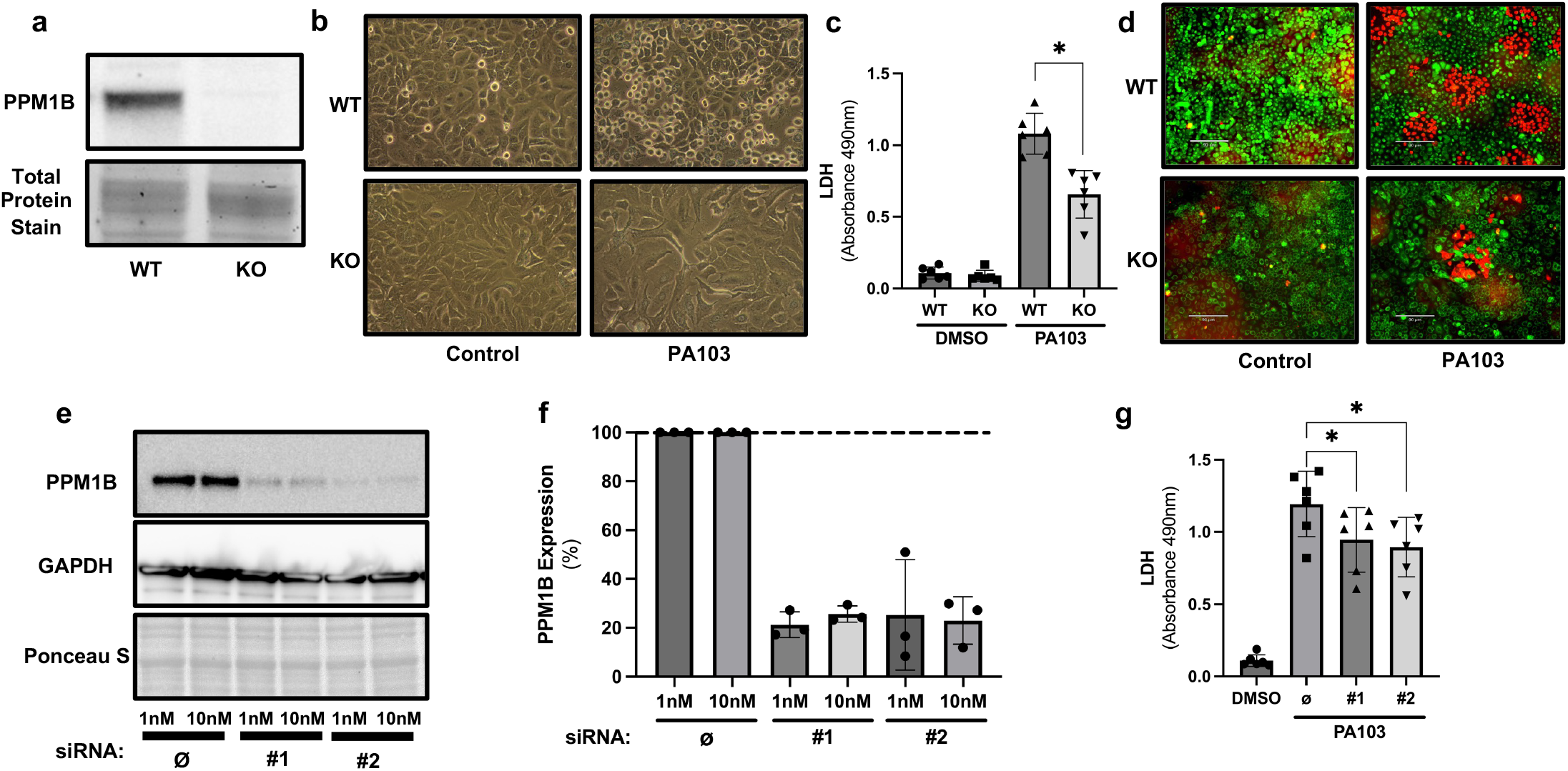
PPM1B promotes cell death during *Pseudomonas aeruginosa* infection. **a**, Western blot of wild-type (WT) and PPM1B knockout (KO) A549 cells. **b**, Representative images of WT cells ± *Pseudomonas aeruginosa* (strain PA103) infection (2 h, 20x magnification). **c**, LDH release from media 2 h post-infection. Data represent mean ± SD of 6 independent experiments; \**P* < 0.05 (one-way ANOVA with Tukey’s post hoc test). **d**, LiveDead staining of cells 2 h post-infection (green = live; red = dead). **e**, Western blot of PPM1B after 48 h treatment with scramble control (Ø) or siRNA (#1, #2) (1 nM or 10 nM) in WT cells. **f**, Quantification of PPM1B knockdown efficiency from three independent experiments (mean ± SD). **g**, LDH release from WT cells treated with Ø or siRNA (#1, #2) 2 h post-PA103 infection (mean ± SD of 6 independent experiments); \**P* < 0.05 (one-way ANOVA with Tukey’s post hoc test).

### A conserved M3 ion is required for catalysis but dispensable for substrate binding

Having defined a functional role of PPM1B in bacterial infection, we next sought to understand the molecular mechanism that enables PPM1B catalytic activity. Crystal structures of human (PPM1A^D146E^, PDB: 6B67, 1.94 Å res), plant (ABI2, PDB: 3UJK; 1.90 Å res), and bacterial (PSTP, PDB: 1TXO, 1.95 Å res) homologs show a third metal ion in the active site that is coordinated by two conserved Asp residues. A sequence alignment indicates PPM1B has these conserved Asp residues at positions Asp151 and Asp243 (Fig. 2a). To investigate the importance of M3, recombinant wild type PPM1B (PPM1B^WT^) and a PPM1B protein with a Asp151Glu mutation (PPM1B^D151E^) was purified (Supplementary Fig. 2) and tested for phosphatase activity. PPM1B^D151E^ did not have detectable phosphatase activity with phosphopeptide substrates and DiFMUP (Fig. 2b-c). PPM1B phosphatase activity was higher with the Promega peptide compared to the IKKβ(pS) phosphopeptide (Fig. 2b). In the plant ortholog PPH1, in vitro activity is enhanced with substrates that have two basic residues preceding the phosphorylation site^15^. Thus, it is likely that PPM1B activity for the Promega peptide is higher due to the preceding “Arg-Arg” basic residues. The binding affinity of M3 is pH sensitive, having the highest affinity at alkaline pH values and the lowest affinity at acidic pH values^13^. If M3 is essential for activity, then a similar pattern in enzymatic activity should be seen across in a pH curve. Indeed, catalytic activity of PPM1B showed peak activity at pH 8.5 with a significant reduction in activity at pH values 7.0 and below (Fig. 2d). Next, a differential scanning fluorimetry assay was performed to assess whether a phosphopeptide, though immune to hydrolysis by PPM1B^D151E^, still binds the mutated catalytic domain. In the presence of an IKKβ(pS) phosphopeptide, PPM1B^D151E^ showed a right shift in melting temperature (T_m_) (average +4.3°C) compared to PPM1B^D151E^ alone (Fig. 2e-f), indicating that the phosphopeptide binds and stabilizes PPM1B^D151E^. Collectively, these results suggest that M3 is critical for catalysis but is dispensable for substrate binding.

**Fig. 2.**
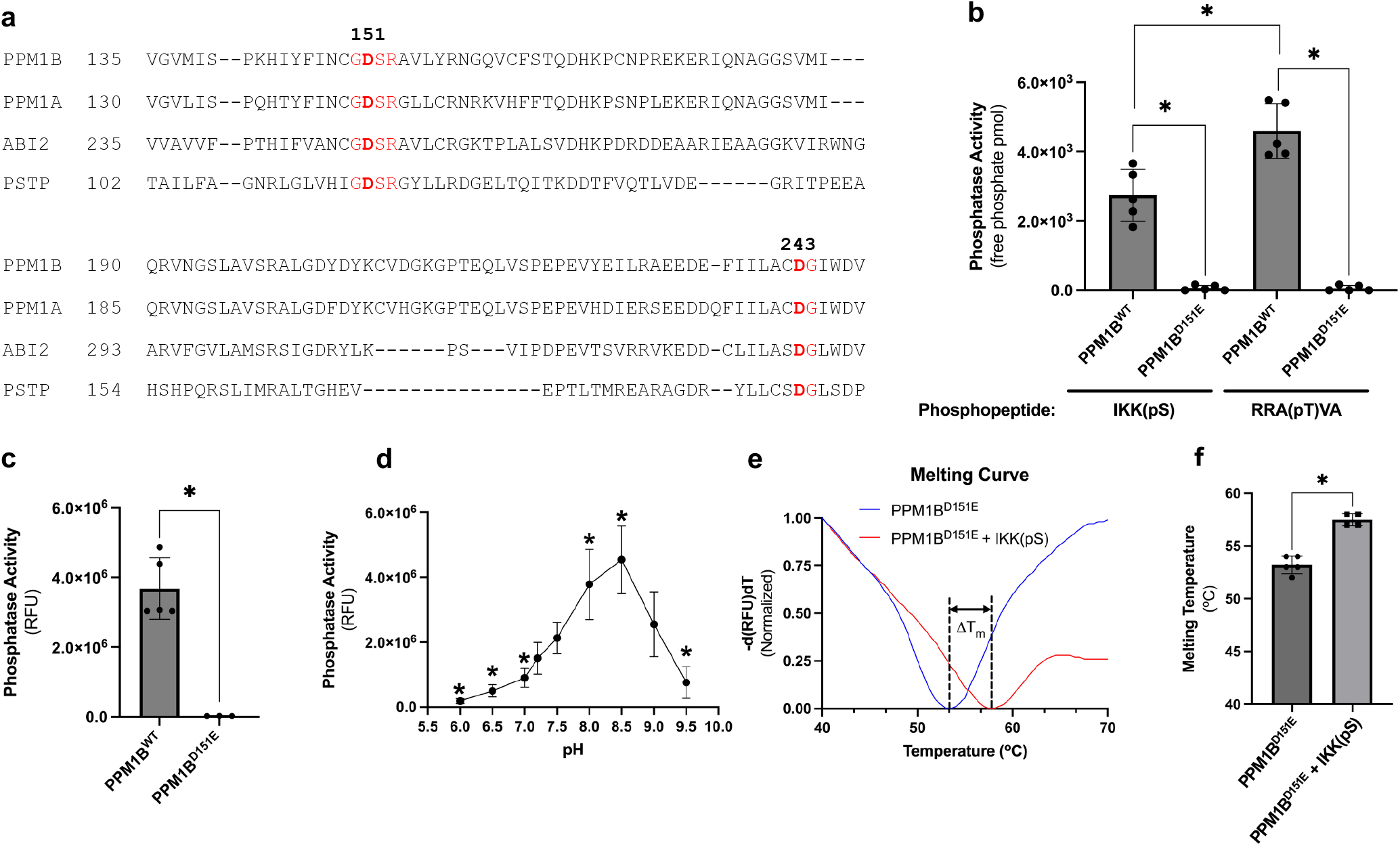
A conserved M3 ion is required for catalysis but dispensable for substrate binding. **a**, MUSCLE sequence alignment of PPM1B with PPM/PP2C homologs from human (PPM1A), plant (ABI2), and bacteria (PSTP). Conserved Asp residues that coordinate the third metal (M3) are highlighted in red. **b, c**, Phosphatase activity of wild-type PPM1B (PPM1B^WT^) and PPM1B with a D151E mutation (PPM1B^D151E^) using the **(b)** phosphopeptide substrates IKKβ(pS) or RRA(pT)VA (100 µM) and **(c)** small molecule substrate DiFMUP (100 µM). Reactions contained 25 nM of enzyme. Data represent mean ± SD of 5 independent experiments; \**P* < 0.05 (one-way ANOVA with Tukey’s post hoc test for **(b)**; two-tailed Student’s t-test for **(c). d**, pH-dependent phosphatase activity of PPM1B^WT^ (25 nM) with DiFMUP (100 µM). Data represent mean ± SD of 8 independent experiments; *P < 0.05 (one-way ANOVA with Tukey’s post hoc test). **e, f**, Differential scanning fluorimetry of PPM1B^D151E^ (5 µM) in the absence or presence of IKKβ(pS) peptide (100 µM). **(e)** Representative melting curves; fluorescence data were normalized on a per-sample basis to a 0-1 range (min-max normalization); vertical dashed lines indicate Tm. **(f)** Quantification of Tm from 4–5 independent experiments; \**P* < 0.05 (two-tailed Student’s t-test).

### A trinuclear metal center positions the substrate phosphate for in-line hydrolysis

To define the role of M3 in PPM1A/1B catalysis, we built a model of PPM1A/1B:phosphoserine complex based on known structures. No crystal structure of wild-type PPM1A or PPM1B containing all three predicted metal ions has been reported. Therefore, crystallization of the PPM1B catalytic domain was attempted to identify their locations in the active site; however, our construct failed to produce diffractable crystals. As an alternative, the crystal structures of PPM1B (PDB: 2P8E; 1.82 Å res.) and PPM1A (PDB: 4RA2; 1.94 Å res.) were superimposed with the plant ortholog ABI2 crystal structure (PDB: 3UJK; 1.90 Å res.) which contains a third Mg^2+^ metal ion in the active site (Fig. 3a). Structural alignment shows PPM1B coordinates M3 with residues Asp151 and Asp243 (with rotation of its carboxylate, Asp243, already M1 bound, should bridge M1 and M3 as in the plant ortholog), and there is an interaction between M3 and Asp60 which bridges M1 and M2 (Fig. 3b). We then placed a phosphoserine in a plausible binding pose within the active site, guided by earlier quantum-based modeling of PP2A-mediated hydrolysis^18^. A cluster model of the PPM1B catalytic site with the phosphoserine and metal ions was constructed using hybrid quantum-based (QM/QM) calculations. In our model, M3 adopts an octahedral coordination geometry, engaging the phosphate group along with two water molecules (W^5^ and W^6^) as ligands (Fig. 3b). All three metals directly bind the phosphoserine’s phosphate moiety, with M1 2.1 Å from phosphoryl oxygen (O^1^), M2 2.2 Å from O^2^, and M3 2.2 Å from O^1^; additionally, Arg33 binds O^3^ (Fig. 3c). M1 and M2 coordinate the bridging hydroxide (W^1^), which functions as the nucleophile for the S_N_2 hydrolase reaction. W^1^ is 3.3 Å from the phosphate center, aligned at an angle of a(W^1^-P-O^γ^) = 164.3° (Fig. 3c). Importantly, O^δ^ of Asp286 hydrogen bonds with W^1^ (2.8 Å) and forms an angle of a(O^δ^-W^1^-P) = 110.3° (Fig. 3c). This geometry positions the nucleophile for in-line attack and suggests that Asp286 constrains the trajectory of W^1^, contributing to orbital steering that promotes optimal overlap with the antibonding orbital of the scissile P-O^γ^ bond and facilitates productive nucleophilic attack. Further analysis of a QM-optimized model also indicates that M3 positions a water molecule (W^5^) to protonate the departing alkoxide (Fig. 4b). This observation suggests that, in addition to contributing to phosphate positioning, M3 plays a direct catalytic role in facilitating the protonation of the leaving group and stabilizing the transition state.

**Fig. 3.**
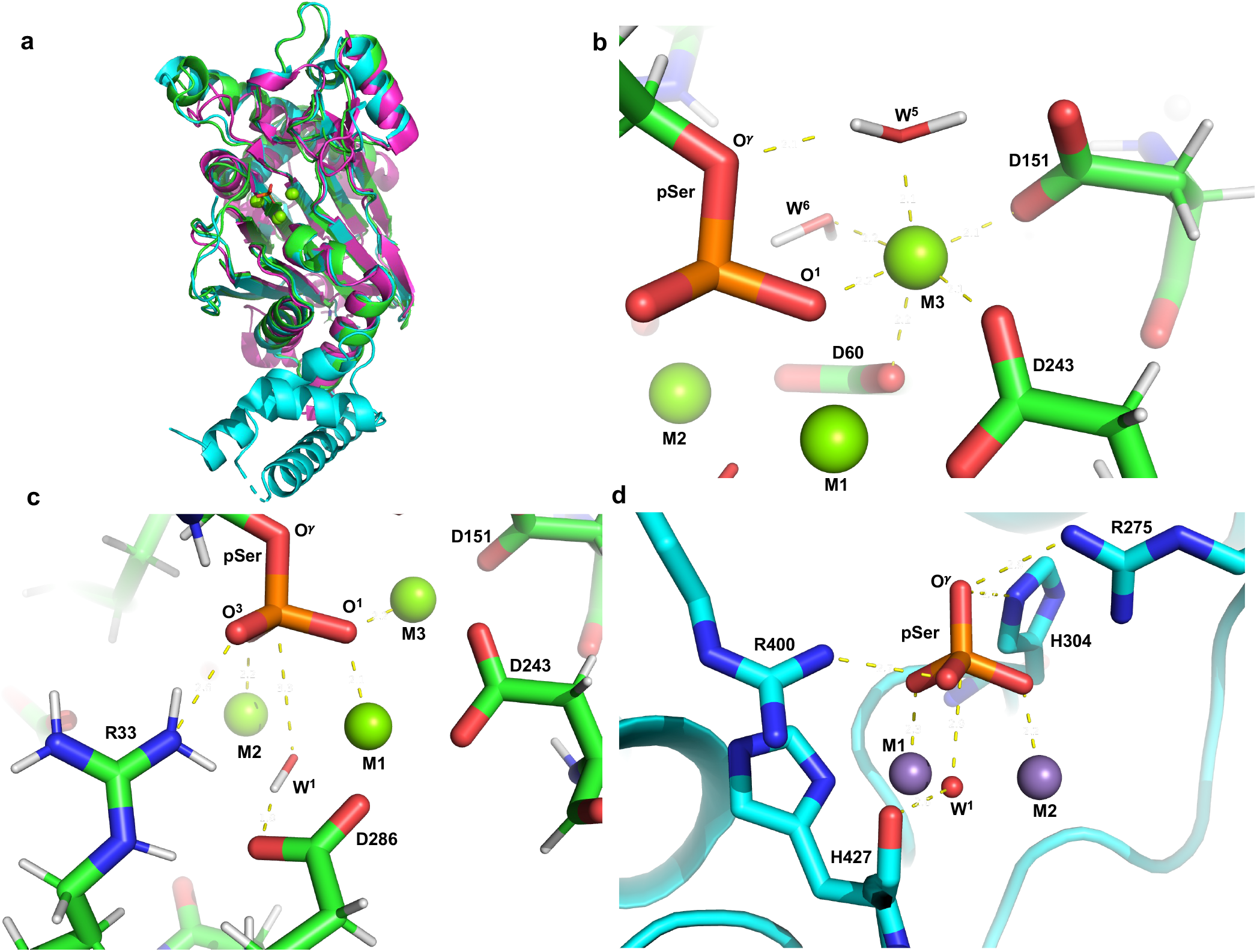
A trinuclear metal center positions the substrate phosphate for in-line hydrolysis. **a**, Structural superimposition of an AMBER optimized model of PPM1B with the crystal structures of human PPM1A (blue; PDE: 4RA2; 1.94 Å res) and the plant ortholog ABI2 (fuchsia; PDE: 3UJK; 1.90 Å res). **b**, QM-optimized model of the PPM1B catalytic site showing octahedral coordination of the third metal ion (M3) with two water molecules (W^5^ and W^6^), a phosphoryl oxygen (O^1^), and residues Asp60, Asp151, and Asp243. W^5^ is positioned for protonation of the leaving alkoxide. **c**, Active-site model of the QM-optimized PPM1B-phosphoserine complex. The trinuclear metal center (M1-M3) and Arg33 coordinate the phosphate to promote in-line nucleophilic attack by the bridging hydroxide (W^1^), with Asp286 constraining W^1^ geometry consistent with orbital steering. **d**, Crystal structure of PP5 (PDB 1S95; 1.60 Å res) showing Arg275 and Arg400 coordinating the phosphate to form the canonical PPP Arg clamp. His427 positions the attacking hydroxide (W^1^) consistent with orbital steering, whereas His304 is positioned to protonate the leaving alkoxide.

**Fig. 4.**
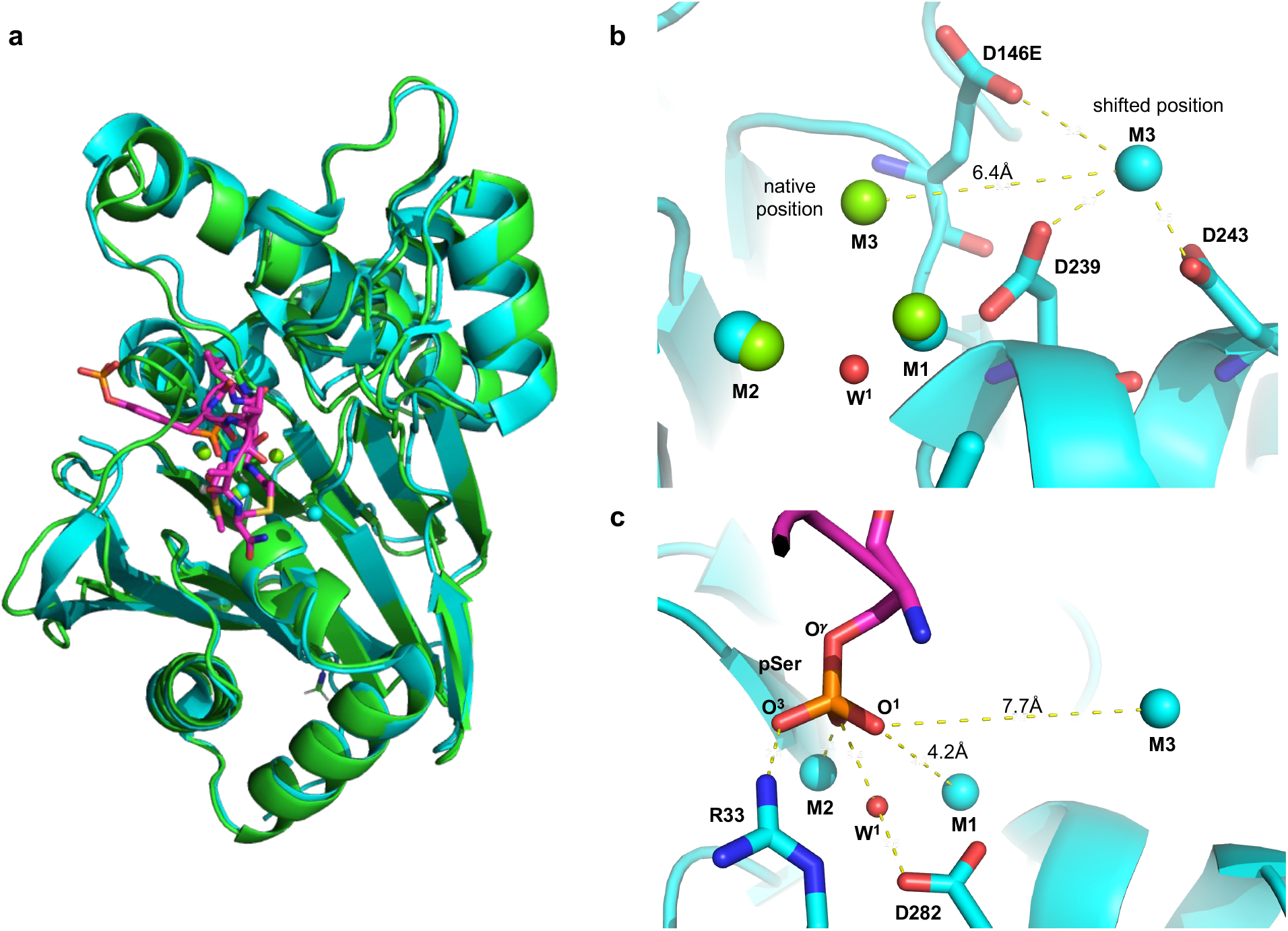
PPM1A^D146E^/PPM1B^D151E^ mutations displace M3 and disrupt catalytic phosphate positioning. **a**, Superimposition of an AMBER-optimized model of the PPM1B (green) and a crystal structure of the inactive PPM1A^D146E^ mutant bound to a cyclic phosphopeptide (blue; PDE: 6B67; 2.20 Å res). **b**, Comparison of residues coordinating the third metal ion (M3) in the PPM1B AMBER-optimized model (green) and the PPM1A^D146E^ crystal structure (blue). **c**, Active site of PPM1A^D146E^ showing the interactions between the substrate phosphate, M2, and Arg33.

To determine whether this mechanism parallels that of PPP phosphatases, we compared the PPM1B model with a crystal structure of PP5 bound with a phosphate (PDB: 1S95; 1.60 Å res.). In PP5, Arg275 and Arg400 form direct contacts with phosphoryl oxygens O^γ^ (2.9 Å) and O^1^ (2.7 Å), respectively, while the two Mn^2+^ ions coordinate O^2^ and O^3^ at distances of 2.2 Å and 2.3 Å (Fig. 3d). This arrangement constitutes the well-characterized Arg clamp that stabilizes and positions the phosphate during catalysis. Notably, the PPM1B trinuclear center establishes a comparable interaction network, in which M3 and Arg33 contact the phosphate in positions analogous to the PPP Arg clamp. These similarities suggest that the third metal in PPM phosphatases functionally substitutes for one component of the Arg clamp, helping to orient the phosphate substrate for nucleophilic attack. Despite these parallels, key mechanistic differences are evident. In PPP phosphatases, orbital steering of the nucleophile has been attributed to backbone elements such as the carbonyl of His427 in PP5 (Fig. 3d). In contrast, our model suggests that PPM1B achieves a similar geometric constraint through Arg286-mediated positioning of the nucleophilic hydroxide. A further divergence is observed in protonation of the leaving-group. In PP5, a conserved His residue (His304) is proposed to protonate the departing alkoxide (Fig. 3d), whereas in PPM phosphatases, this role is fulfilled by a M3-bound water molecule. Together, these findings highlight a convergent catalytic strategy in which both enzyme families achieve efficient phosphoryl transfer, while employing fundamentally different active-site architectures to control nucleophilic alignment and leaving-group stabilization.

### PPM1A^D146E^/PPM1B^D151E^ mutations displace M3 and disrupt catalytic phosphate positioning

To rationalize the loss of catalytic activity in the PPM1B^D151E^ mutation, we compared our wild type PPM1B model with the co-crystal structure of the inactive PPM1A^D146E^ mutant complexed with a cyclic phosphopeptide (PDB: 6B67; 2.20 Å res) (Fig. 4a). Side-by-side comparison reveals that the Asp146Glu mutation shifts M3 6.4 Å away from the expected native position (Fig. 4b), resulting in no interaction between M3 and the cyclic peptide’s phosphate. The shifted position introduces a new interaction between M3 and Asp243 in PPM1A^D146E^ (i.e., Asp247 in PPM1B) (Fig. 4b) which is not seen in our structural model of the wild type PPM1A/1B (Fig. 3b). Prior mutagenesis studies show Asp243 in PPM1A does not participate in enzymatic activity but helps PPM1A retain binding affinity to M3 when Asp146 is mutated to an Ala^13^, suggesting that the interaction stabilizes M3 in the shifted location. In the PPM1A^D146E^ mutant, M2 directly binds one phosphoryl oxygen, but the phosphate group is not directly liganded to M1 and M3, with the nearest phosphoryl oxygen at a distance of 4.2 Å away from M1 and 7.7 Å from M3 (Fig. 4c). While Arg33 also binds one of the peptide’s phosphate oxygens, Arg33 likely arrests the phosphate group in a nonreactive position due to lack of counter pull from a natively positioned M3. Consequently, the cyclic peptide’s phosphate is misaligned for the nucleophilic attack by W^1^ with a W^1^-P distance of 3.4 Å and a(W^1^-P-O^γ^) = 146.7° (Fig. 4c). Furthermore, loss of direct M1 and M3 phosphate binding in the PPM1A^D146E^ mutant likely results in insufficient withdrawal of electron density from the central P atom to support the S_N_2 reaction. Overall, these findings suggest that PPM1A/1B has a third metal that directly binds O^1^ in the active site, which is critical for both positioning the phosphate and for drawing sufficient electron density out of the central phosphorous atom so that the nucleophilic attack by W^1^ can occur.

## Discussion

While previous studies indicated M3 is essential for activity^13–17^, the molecular basis for its contribution to phosphate positioning and catalysis has not been defined. Our data and structural modeling provide the first mechanistic explanation for how the PPM family utilizes a unique trinuclear metal architecture to directly orchestrate phosphate positioning and catalysis. Notably, the position of Arg33 is variable between PPM crystal structures, and it has been previously reported to have the largest conformational variation during molecular dynamics simulation^14^. Thus, it is likely that Arg33 is a flexible residue that helps catch and guide the substrate to the three metals. Structural modeling supports a mechanism in which M3, in conjunction with Arg33, directly coordinates a phosphoryl oxygen, positioning the phosphate group for in-line nucleophilic attack by the bridging hydroxide coordinated by M1 and M2. This configuration simultaneously orients the substrate and withdraws electron density from the central phosphorus atom, creating conditions favorable for the S_N_2 hydrolase reaction. In line with prior studies^14,15,17,19^, our model further supports a catalytic role for M3 beyond substrate positioning, in which an M3-bound water is optimally positioned to protonate the departing alkoxide, thereby facilitating leaving-group departure and stabilizing the transition state.

A striking feature of PPM phosphatase catalysis is its convergence with the catalytic strategy used by PPP phosphatases, despite fundamentally different active-site architectures. In PPP enzymes, substrate positioning and transition state stabilization are mediated by an Arg clamp and protein-based orbital steering elements, with a conserved histidine serving as the proton donor to the leaving group^9–12^. In contrast, PPM phosphatases redistribute these functions across a trinuclear metal center, where M3 substitutes for a component of the Arg clamp and also enables leaving-group protonation via a coordinated water molecule. These findings reveal that two evolutionarily distinct phosphatase families have converged on equivalent catalytic solutions for phosphate positioning and orbital alignment, using either basic residues or a metal ion to achieve efficient phosphoryl transfer. Moreover, M3 coordinating residues are conserved across bacteria, plant, insect, fungi, and mammal PPM/PP2C phosphatases^20^, indicating that it is an ancient mechanism that developed early in evolution and is broadly utilized by members of the PPM/ PP2C family.

Despite the clear mechanistic importance and evolutionary conservation of the M3 site, it remains unclear how M3-dependent catalysis is supported in cells, given that millimolar concentrations of Mg^2+^ or Mn^2+^ are required for M3 occupancy in vitro, whereas cytosolic Mg^2+^/Mn^2+^ levels are estimated to be 0.5-1.0 mM^21–23^. Recent reports indicate that PPM1B can undergo liquid-liquid phase separation (LLPS) in cells^24^. More generally, LLPS has been proposed as a mechanism that can locally modulate the concentrations of proteins and associated small molecules, including ions^25^. In a simple two-compartment model, we find that partitioning PPM1B into a LLPS condensate is predicted to increase local Mg^2+^ concentrations to levels compatible with M3 occupancy (Supplementary Fig. 3 and Supplementary Table 1). Although indirect, these observations raise the possibility that cellular compartmentalization could influence metal availability at the M3 site. Future studies will be required to determine whether and how such effects may contribute to PPM1B regulation in vivo.

Finally, we observed that PPM1B is linked to cell death during bacterial challenge. Although the substrates responsible for this effect remain undefined, PPM1B targets multiple components of innate immune and cell death pathways^1–8^, and pathogen-encoded effectors can redirect PPM1B activity to subvert host immunity^5,26,27^. Consistent with a broader role for this phosphatase family, other PPM members, including PPM1A and PPM1G, have been incriminated in the pathology of bacterial and viral infections^28–32^. Notably, the catalytically inactive Asp151Glu mutant of PPM1B, as well as equivalent substitutions in related homologs^14^, retains substrate binding while abolishing enzymatic activity. These features suggest that such mutants could be used to stabilize transient enzyme-substrate complexes in cells, enabling systematic identification of physiological substrates and help define how microbial effectors reprogram PPM1B and related phosphatases during infection.

In summary, our findings reveal PPM1B promotes cell death during bacterial infection, and PPM phosphatases utilize a trinuclear metal center to enable catalysis. Stable trinuclear metal centers are exceptionally rare catalytic architectures in the human proteome, occurring in only a small number of enzyme families^13,14,33^. Our findings indicate that PPM phosphatases represent a previously underappreciated example of this rare catalytic strategy, using a third metal ion to directly control substrate positioning during phosphoryl hydrolysis. Given the absence of clinically used small-molecule inhibitors for PPMs and phosphatases in general, this trinuclear metal architecture represents a promising target for structure-based drug design for immune and infectious diseases.

## Materials and Methods

### Generation of PPM1B knockout cells

Synthetic, chemically modified single guide RNAs (sgRNAs) and cas9 nuclease (Cas9) was ordered from Synthego. Briefly, electroporation of Cas9 protein and three different sgRNAs that target exon 2 in the PPM1B gene was done on A549 cells. After 48-hours, treated cells were subjected to single cell cloning and DNA was isolated and sequenced. Positive hits were single cell cloned a second time and isolated DNA was sequenced to confirm successful insertion of a premature stop codon. Knockout cells were further verified with quantitative PCR and western blotting.

### siRNA knockdown of PPM1B in cells

Scramble control (Ø) and knockdown siRNA (#1 and #2) were ordered from ThermoFisher Scientific. Cells were seeded at 4.0x10^5^ cells per well in a 6-well plate. The following day, cells were rinsed with PBS and transfected with 10 nM of siRNA-lipofectamine RNAiMAX complex in antibiotic free DMEM media supplemented with FBS, pyruvate and non-essential amino acids. Knockdown efficiency was assessed with western blot and infection experiments were performed 48-hours after siRNA treatment.

### Western blotting

Cells were lysed in RIPA buffer supplemented with protease and phosphatase inhibitor cocktails. Lysates were centrifuged at 14,000 rpm for 10 min at 4°C, and the supernatant was collected as the protein extract. Samples were mixed with Laemmli buffer, heated in a water bath at 100°C for 10 min, and resolved on a 12% gel (Mini-PROTEAN TGX Stain-Free Protein Gel; Bio-Rad). Total protein was visualized using stain-free imaging. Proteins were then transferred to a nitrocellulose membrane and blocked in 5% non-fat milk in TBST. Membranes were incubated with primary antibodies against PPM1B (1:2,000; ProteinTech; 13193-1-AP) or GAPDH (1:1,000; Cell Signaling Technology; #2118), followed by incubation with HRP-conjugated goat anti-rabbit secondary antibody (1:10,000; Abcam) for 1 hour at 4°C. Blots were developed using SuperSignal West Femto Maximum Sensitivity Substrate (ThermoFisher Scientific) and imaged. Band intensities were quantified using ImageJ.

### *Pseudomonas aeruginosa* infection assay

Infections were performed as previously described^34,35^. Cells were seeded at 4.0x10^5^ cells per well in a 6-well plate and grown for three days in DMEM media supplemented with FBS, pyruvate and non-essential amino acids. *Pseudomonas aeruginosa* (strain PA103) was grown on Vogel-Bonner agar plates overnight at 37°C. The day of the infection, PA103 was suspended in PBS until an OD_540_ of 0.250 was reached (ThermoFisher Scientific, AquaMat Plus UV-VIS), which has previously been shown to equal 2.0x10^8^ colony forming units (CFUs) per mL^34,35^. Cells were then infected at a 20:1 multiplicity of infection for 2 hours in HBSS media. Afterwards, LDH from the media was measured, and LiveDead assay was performed.

### Protein purification

Plasmids for recombinant human PPM1B and PPM1B^D151E^ proteins were ordered from GeneArt, and Gateway cloning was done to insert plasmids into a His-MBP expression vector (Addgene: plasmid #11085)^36^. Proteins were transfected into *E. coli* Rosetta (DEL3) cells (Millipore Sigma) and selected for with ampicillin agar plates (100 µg/mL) overnight. Single colonies were isolated and grown in a starter culture containing LB broth with ampicillin (100 µg/mL) over night. The following day, an aliquot of the starter culture was grown in 5 L of LB broth until an OD_600_ of 0.6 – 1.0 was reached, and protein expression was then induced with 0.5 mM of IPTG overnight at 30°C. Bacteria were pelleted and frozen in -80°C. During purification, bacteria pellets were lysed with B-Per lysis buffer (4 mL B-per buffer/ 1 g bacteria, 200 µg lysozyme/ mL buffer, 10U DNase I/ mL buffer, 2 mM MgCl_2_, and 1 EDTA-free protease inhibitor tablet/ 8 mL) (ThermoFisher Scientific) for 1.5 hours at room temperature, followed by another 1.5 hours of digestion on ice, and centrifugation at 33,000 x g for 40 minutes at 4°C. The supernatant was collected, sonicated, and equilibrated with Ni binding buffer (20 mM Tris base, 500 mM NaCl, 1 mM MnCl_2_, 5% glycerol, and 20 mM imidazole). Recombinant protein was then captured with HisTrap (Cytiva) affinity chromatography and eluted with Ni elution buffer (20 mM Tris base, 500 mM NaCl, 1 mM MnCl_2_, 5% glycerol, and 500 mM imidazole). Protein was supplemented with approximately 500 mg of TEV protease and dialyzed with dialysis buffer (40 mM Tris base, 1 mM MnCl_2_, 5% glycerol, 0.5 mM EDTA) for 8 hours at room temperature, followed by another dialysis for an additional 10 hours at 4°C. Cleaved His-MBP tag was removed by running the protein through another HisTrap or MBPTrap affinity column and untagged protein was then further purified with size exclusion chromatography (HiLoad 16/600 Superdex 200 pg, Cytiva) equilibrated with column buffer (20 mM HEPES, 1mM MnCl_2_, 5% glycerol, and 300 mM NaCl). Finally, protein was diafiltrated into storage buffer (20 mM HEPES, 1mM MnCl_2_, 5% glycerol), concentrated, and flash-frozen in liquid nitrogen before being stored in aliquots at -80°C. Purified phosphatases were confirmed with enzymatic activity assays and western blotting.

### Phosphatase activity assays

For activity assays, proteins were stored at -20°C in a 50% storage buffer: 50% glycerol mixture, and assays were performed within 1-week of thawing a protein aliquot. Phosphopeptide activity assays were performed with the Promega Serine/Threonine Phosphatase Assay System (V2460). A phosphopeptide derived from IKKβ(pS) (residues 175-184, QG(pS)LCT(pS)FVG) was ordered from CPC Scientific and used in the assay. A synthetic phosphopeptide (RRA(pT)VA) from Promega was also used as an additional substrate, and activity assays were performed for 30 minutes at 30°C in activity assay buffer (20 mM 7.5 pH HEPES, 1 mg mL^-1^ BSA, 5 mM MnCl_2_, 1 mM DTT, 0.2 mM EGTA). Phosphatase activity with 6,8-difluoro-4-methylumbelliferyl phosphate (DiFMUP) (ThermoFisher Scientific) was monitored by measuring fluorescence (ex = 358 nm and em = 450 nm) for 30 minutes at 30°C in DiFMUP buffer (30 mM 7.5 pH HEPES, 1 mg mL^-1^ BSA, 1 mM NaAscorbate, 0.1% Triton X-100, 1 mM DTT, and 1 mM MnCl_2_) as previously described^37,38^. For the pH curve, a three-way buffer (50 mM acetic acid, 50 mM MES, 100 mM Tris, 1 mg mL^-1^ BSA, 1 mM NaAscorbate, 0.1% Triton X-100, 1 mM DTT, and 1 mM MnCl_2_) was used to maintain a consistent ionic strength across all pH values tested.

### Differential scanning fluorimetry

PPM1B^D151E^ was labeled with SYPRO Orange dye (ThermoFisher Scientific) and subjected to thermal denaturation using a Bio-Rad CFX Connect qPCR instrument in assay buffer (pH 7.0, 30 mM HEPES, 1 mM sodium ascorbate, 5 mM MnCl_2_, 0.01% Triton X-100, 5 µM PPM1B^D151E^ protein) in the presence or absence of the phosphopeptide IKKβ(pS) (100 µM). Fluorescence data were normalized on a per-sample basis by scaling to the pre- and post-transition baselines (F_min_ and F_max_) to a 0-1 range, thereby accounting for differences in signal intensity between conditions. The melting temperature (T_m_) was then calculated and compared between groups.

### Quantum based modeling of PPM1A/B structures

Our model PPM1B:phosphoserine complex was prepared for AMBER24 molecular mechanics (MM) optimization under the ff14SBonlysc force field^39^. First, we used AlphaFold 3 to generate missing residues in PDB: 2P8E (1-3; 87-97; 125-127)^40^, completing the sequence through Phe295. An N-methyl termination (NME) was added, and His162 and His169 were protonated (HIP). M1, M2, and M3 were assigned as Mg^2+^ ions. The bridging hydroxide ion W^1^ was assigned gaff parameters with custom charges (O: o, -1.205; H: ho, +0.205). Four explicit waters associated with the tri-metal system as found in PDB: 3UJL were included in our model: two waters (W^2^, W^3^) to complete the octahedral center at M2; one water (W^4^) to potentially complete the octahedral center at M1; and one water (W^6^) coordinated with M3 that is also H-bonded to His162. Additionally, one water (W^5^) was introduced to give M3 potentially a 5^th^ or 6^th^ coordination, depending upon the orientations of Asp60 and Asp243. All waters were assigned TIP3P parameters. The phosphoserine ligand (SEP; phosaa14SB) was terminated with acetyl (ACE) and N-methyl (NME) groups. The system was solvated with TIP3P waters in an octahedral box with a 10 Å buffer; ions were added to balance the system’s charge and to mimic 0.150 M NaCl. A 10,000-step optimization was performed in which the metals, the O atom of the bridge hydroxide, and the central P atom of the phosphoserine’s phosphate were restrained. After the MM optimization, we used AMBER to extract a zero-charge cluster model^39^, composed of the phosphoserine ligand, the tri-metal system, and 14 residues of the catalytic site. At breaks in the backbone, residues were terminated by AMBER as aldehydes and unprotonated amines. We then carried out a quantum-based hybrid optimization on the cluster model using Gaussian16 (ONIOM) in implicit water (SCRF)^41^. To the high-level region, we applied the wB97XD density functional and cc-pvtz basis set; for the low level-region, which was frozen during the optimization, we used the semi-empirical PM7 method.

## Acknowledgments

This work was supported by the National Science Foundation (NSF-2450845) and the American Cancer Society (RSG-25-1412630-01-DMC), both to A.P. Additional support was provided by start-up funds from the Mitchell Cancer Institute (to A.P.) and seed funding from the University of South Alabama Intramural Grants Program (to A.P. and M.R.S.). This work was also supported by the National Institutes of Health (R01HL160988 to J.Y.L.). Computational resources and technical support were provided in part by the Alabama Supercomputer Authority.

---

**Supplementary Fig. 1.**
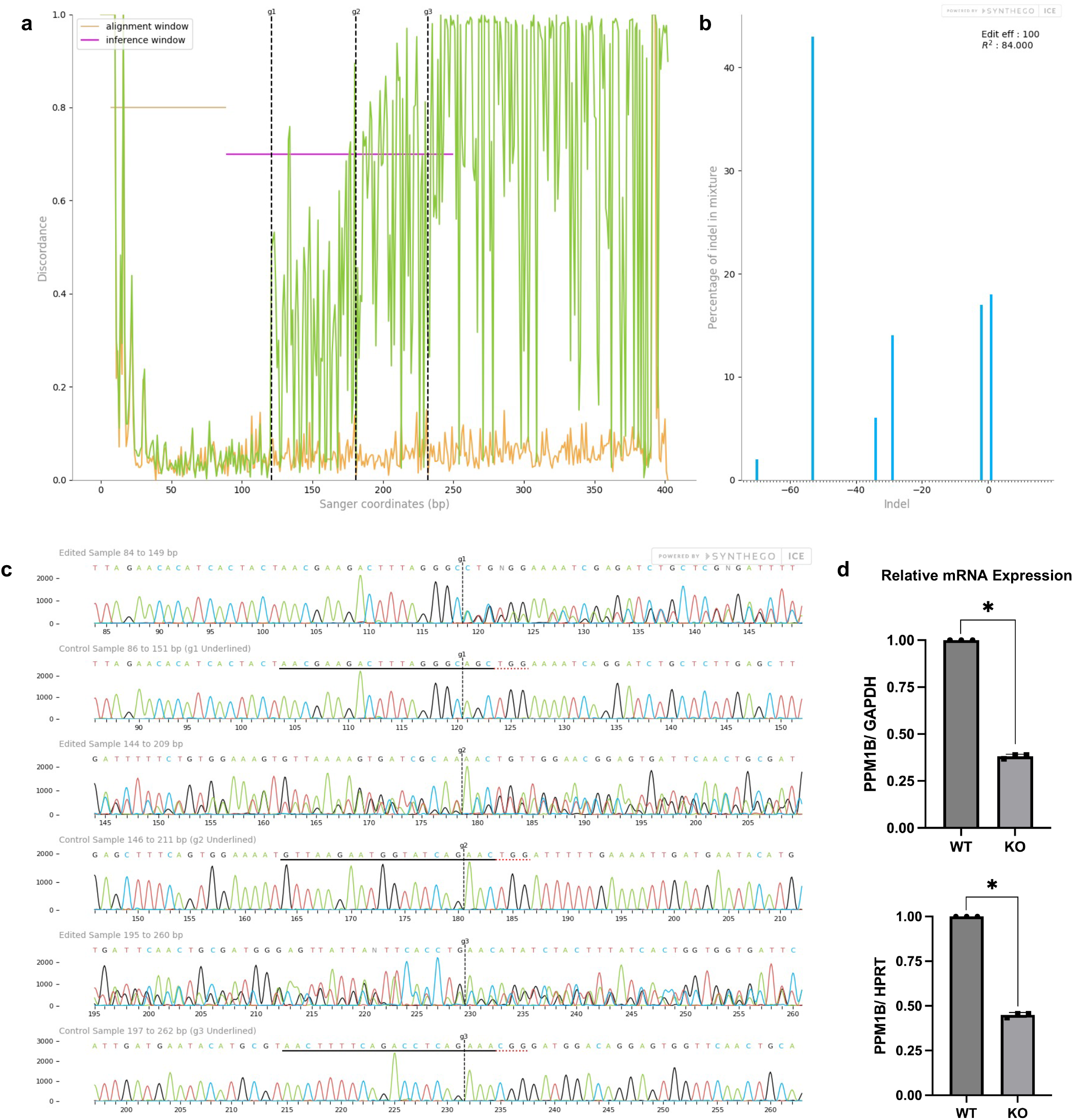
Validation of PPM1B knockout in A549 cells. **a**, Discordance plot comparing Sanger sequences of WT (orange) and CRISPR KO (green) cells. Dotted black lines indicate guide RNA cut sites (g1–g3). **b**, Indel plot showing frequency and size distribution of insertions/deletions in KO cells. **c**, Representative Sanger traces of WT and KO cells at guide RNA target sites; black underline = guide sequence; dotted red line = PAM; dotted vertical line = cut site. **d**, Quantitative PCR showing PPM1B mRNA levels normalized to GAPDH and HPRT.

**Supplementary Fig. 2.**
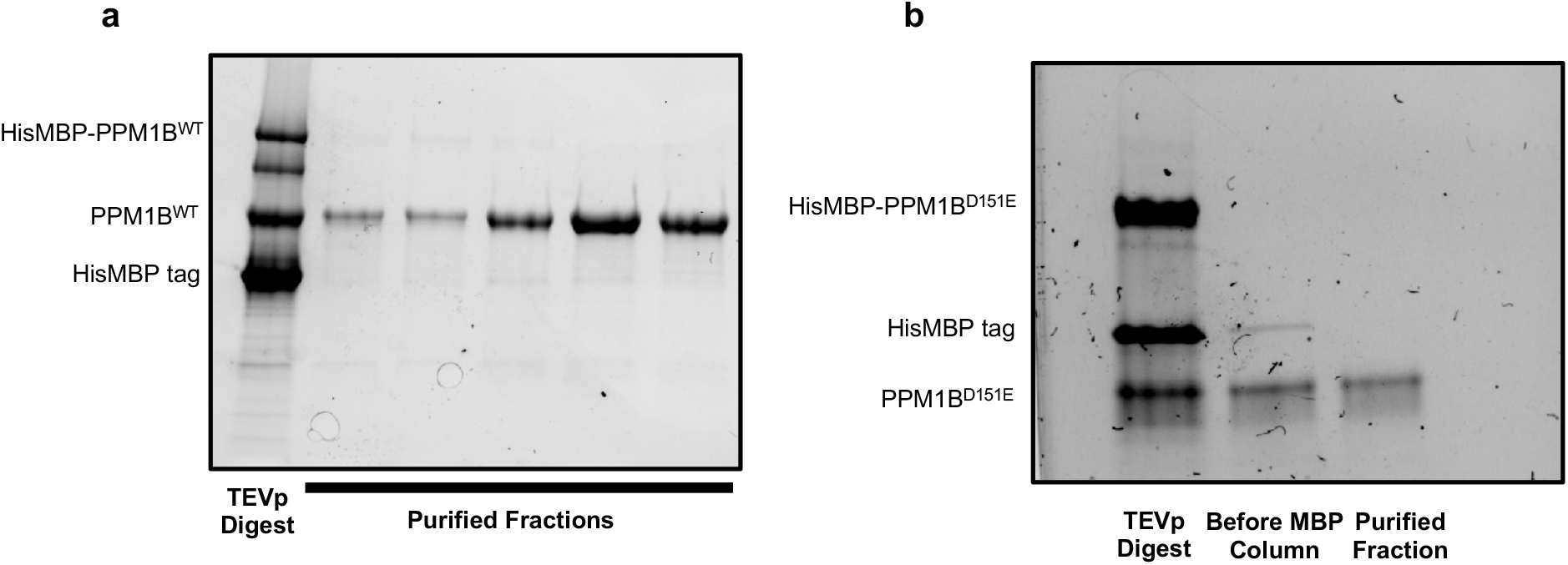
Purification of recombinant human PPM1B proteins. **a, b**, Gels showing TEV protease (TEVp) digestion and final purified fractions of (**a**) wild type PPM1B (PPM1B^WT^) and (**b**) PPM1B with a D151E mutation (PPM1B^D151E^) after size-exclusion chromatography.

**Supplementary Fig. 3.**
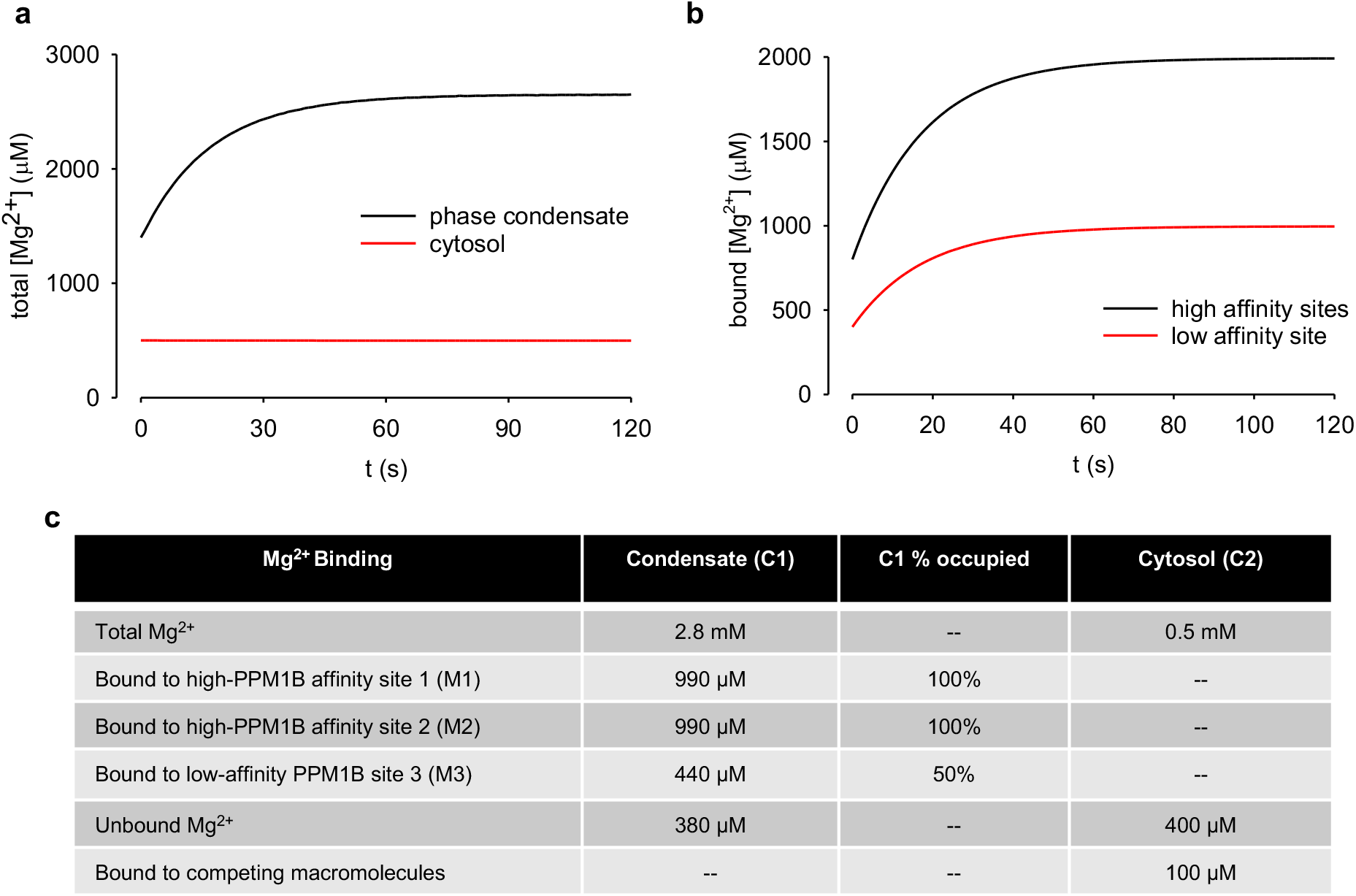
Simulation of Mg^2+^ distribution in cytosol and liquid-liquid phase condensates. **a**, Modeled distributions of total Mg^2+^ in the cytosol (red) and phase condensates (black). Kinetics are dominated by diffusion constraints at the condensate-cytosol interface. **b**, Mg^2+^ binding kinetics to low- and high-affinity PPM1B sites within the phase condensate. High-affinity sites bind faster than cytosolic flux. **c**, Occupancy of Mg^2+^ at high-(M1 and M2) and low-affinity (M3) sites in a 100 nm sized condensate with 1 mM PPM1B. High-affinity sites are fully occupied, low-affinity sites reach ∼50% occupancy under modeled conditions.

**Supplementary Table 1.** Parameter definitions, values, and initial conditions used to simulate Mg^2+^ sequestration into the PPM1B phase condensate.

| variable | description | value | initial condition | steady state |
| --- | --- | --- | --- | --- |
| V1 | volume of compartment 1, the phase condensate | 0.002 pL | – | – |
| V2 | volume of compartment 2, the bulk cytosol | 2.0 pL | – | – |
| n1 | total $[Mg^{2+}]$ in the phase condensate | – | $1.40 \times 10^{-3} \text{ M}$ | $2.65 \times 10^{-3} \text{ M}$ |
| n2 | total $[Mg^{2+}]$ in the cytosol | – | $0.50 \times 10^{-3} \text{ M}$ | $0.50 \times 10^{-3} \text{ M}$ |
| n3 | [PPM1B] without $Mg^{2+}$ bound to high affinity binding site 1 | – | $0.70 \times 10^{-3} \text{ M}$ | $0.63 \times 10^{-3} \text{ M}$ |
| n4 | [PPM1B] with $Mg^{2+}$ bound to high affinity binding site 1 | – | $0.30 \times 10^{-3} \text{ M}$ | $0.37 \times 10^{-3} \text{ M}$ |
| n5 | [PPM1B] without $Mg^{2+}$ bound to high affinity binding site 2 | – | $0.60 \times 10^{-3} \text{ M}$ | $0.00 \times 10^{-3} \text{ M}$ |
| n6 | [PPM1B] with $Mg^{2+}$ bound to high affinity binding site 2 | – | $0.40 \times 10^{-3} \text{ M}$ | $1.00 \times 10^{-3} \text{ M}$ |
| n7 | [PPM1B] without $Mg^{2+}$ bound to low affinity binding site | – | $0.60 \times 10^{-3} \text{ M}$ | $0.00 \times 10^{-3} \text{ M}$ |
| n8 | [PPM1B] with $Mg^{2+}$ bound to low affinity binding | – | $0.40 \times 10^{-3} \text{ M}$ | $1.00 \times 10^{-3} \text{ M}$ |
| n9 | [high affinity cytosolic buffer] without $Mg^{2+}$ bound | – | $0.10 \times 10^{-3} \text{ M}$ | $0.00 \times 10^{-3} \text{ M}$ |
| n10 | [high affinity cytosolic buffer] with $Mg^{2+}$ bound | – | $0.90 \times 10^{-3} \text{ M}$ | $0.10 \times 10^{-3} \text{ M}$ |
| n11 | [low affinity cytosolic buffer] without $Mg^{2+}$ bound | – | $1.90 \times 10^{-3} \text{ M}$ | $1.89 \times 10^{-3} \text{ M}$ |
| n12 | [low affinity cytosolic buffer] with $Mg^{2+}$ bound | – | $0.10 \times 10^{-3} \text{ M}$ | $0.11 \times 10^{-3} \text{ M}$ |
| k1 | $Mg^{2+}$ flux coefficient from the phase condensate into the cytosol | $0.01 \times 10^{-6} \text{ L/s}$ | – | – |
| k2 | $Mg^{2+}$ flux coefficient from the cytosol into the phase condensate | $0.01 \times 10^{-6} \text{ L/s}$ | – | – |
| $k_{on1}$ | rate constant for $Mg^{2+}$ binding to low affinity PPM1B binding sites | $200 \text{ M}^{-1}\text{s}^{-1}$ | – | – |
| $k_{off1}$ | off rate for the low affinity PPM1B binding site | $0.1 \text{ s}^{-1}$ | – | – |
| $k_{on2}$ | rate constant for $Mg^{2+}$ binding to high affinity PPM1B binding sites | $200 \text{ M}^{-1}\text{s}^{-1}$ | – | – |
| $k_{off2}$ | off rate for high affinity PPM1B binding sites | $2 \times 10^{-4}$ | – | – |
| $k_{on3}$ | rate constant for binding to high affinity cytosolic $Mg^{2+}$ binding sites | $200 \text{ M}^{-1}\text{s}^{-1}$ | – | – |
| $k_{off3}$ | off rate for the high affinity cytosolic $Mg^{2+}$ binding sites | $2 \times 10^{-4}$ | – | – |
| $k_{on4}$ | rate constant for $Mg^{2+}$ binding to low affinity cytosolic binding sites | 200 | – | – |
| $k_{off4}$ | off rate for $Mg^{2+}$ binding to low affinity cytosolic sites | $1 \text{ s}^{-1}$ | – | – |

